# Comparison of dipsogenic responses of adult offspring as a function of different perinatal programming models

**DOI:** 10.1101/2021.10.14.464417

**Authors:** FM Dadam, JL Amigone, L Vivas, AF Macchione

**Affiliations:** Instituto de Investigación Médica Mercedes y Martín Ferreyra, INIMEC-CONICET-Universidad Nacional de Córdoba, Córdoba, Argentina; Instituto de Investigaciones Psicológicas, IIPsi-CONICET-Universidad Nacional de Córdoba, Córdoba, Argentina; Facultad de Ciencias Exactas, Físicas y Naturales, Universidad Nacional de Córdoba, Córdoba, Argentina; Sección de Bioquímica Clínica, Hospital Privado, Córdoba, Argentina

**Keywords:** pre/postnatal imprinting, partial aortic ligation, chronic over-activation of RAS, hypovolemic thirst, absolute dehydration, relative dehydration, dorsal raphe nucleus

## Abstract

The perinatal environment interacts with the genotype of the developing organism resulting in a unique phenotype through a developmental or perinatal programming phenomenon. However, it remains unclear how this phenomenon differentially affects particular targets expressing specific drinking responses depending on the perinatal conditions. The main goal of the present study was to compare the dipsogenic responses induced by different thirst models as a function of two perinatal manipulation models, defined by the maternal free access to hypertonic sodium solution and a partial aortic ligation (PAL-W/Na) or a sham-ligation (Sham-W/Na). The programmed adult offspring of both perinatal manipulated models responded similarly when was challenged by overnight water dehydration or after a sodium depletion showing a reduced water intake in comparison to the non-programmed animals. However, when animals were evaluated after a body sodium overload, only adult Sham-W/Na offspring showed drinking differences compared to PAL and control offspring. By analyzing the central neurobiological substrates involved, a significant increase in the number of Fos + cells was found after sodium depletion in the subfornical organ of both programmed groups and an increase in the number of Fos + cells in the dorsal raphe nucleus was only observed in adult depleted PAL-W/Na. Our results suggest that perinatal programming is a phenomenon that differentially affects particular targets which induce specific dipsogenic responses depending on matching between perinatal programming conditions and the osmotic challenge in the latter environment. Probably, each programmed-drinking phenotype has a particular set point to elicit specific repertoires of mechanisms to reestablish fluid balance.

## 1. Introduction

Gestational characteristics modify the intrauterine environment and differentially program the developing organism. Studies carried out in the last decades of the 20th century have provided evidence of the important role of the environment when it acts during critical periods of the intrauterine and/or early postnatal life (Lejarraga, 2019). Hence, a widely recognized hypothesis that considers the environmental influence on the new individual and its long-term effects is the “fetal origin hypothesis”, postulated by Barker in the 1990s (Barker, 2002; Barker et al., 2002; de Boo & Harding, 2006). These environment-genotype interactions over a sensitive period result in an unequivocal phenotype expression induced by particular plasticity processes called perinatal programming or developmental plasticity (Mecawi et al., 2015). This phenomenon affects a wide variety of systems during development, including the hydroelectrolyte homeostatic system that regulates thirst, sodium appetite, renal excretion, blood pressure, etc., through neural and endocrine mechanisms.

Different pre- and/or postnatal persistent endocrine and osmotic conditions during ontogenetic sensitive periods can modify regulation mechanisms of offspring’s fluid intake, even in adulthood (Mecawi et al., 2015). Longitudinal studies reveal permanent alterations to sodium-preference, spontaneous sodium intake, and depletion-induced sodium intake in adult offspring gestated in a perinatal environment characterized by conditions of sodium overload or sodium depletion (Contreras & Kosten, 1983; Contreras & Ryan, 1990; Crystal & Bernstein, 1995, 1998; Curtis et al., 2004; Galaverna et al., 1995; Shirazki et al., 2007).

In animal preclinical studies, we have demonstrated that the mere ad-libitum access to a hypertonic sodium solution during pregnancy and lactation (i.e. voluntary maternal intake) is sufficient to induce an offspring’s osmoregulatory programming that persist into adulthood (Macchione et al., 2012, 2015). We tested the long-term effects of this perinatal programming paradigm in a hypovolemic thirst model (Macchione et al., 2012). The maternal ad-libitum sodium intake was observed to induce a significant reduction of sodium and water intake stimulated by Furosemide and a low sodium diet treatment in adult offspring. These alterations in fluid intake were also linked to modifications in the brain cell activity patterns of central areas involved in the hydroelectrolyte balance regulation. The programmed adult offspring exhibited an increased number of Fos immunoreactive cells in the subfornical organ (SFO) and supraoptic nucleus (SON) in comparison with non-programmed animals. These results were the first to demonstrate that the maternal voluntary intake of a hypertonic solution during pregnancy and lactation induces an effective and endurance offspring programming. Besides, this investigation also showed that the behavioral effects may be explained by functional modifications at the central nervous system.

In this perinatal programming model, we have also studied whether there was any programming effect in adult offspring when employing another osmotic challenge, such as the intravenous infusion of hypertonic sodium solution (Macchione et al., 2015). This osmotic challenge is an opposite stimulus in some aspects to the previous ones, because instead of body sodium and water depletion, a body sodium overload is induced, eliciting thirst and activating the peripheral and central pathways triggered by osmo-sensors stimulated by the increase in the CSF and plasma osmolality (McKinley & Johnson, 2004). Once again, alterations in both behavioral and underlying neurobiological substrates were identified. In this case, opposite programming effects were observed in the programmed offspring, i.e., a significant increase in water intake and a decrease in the number of Fos immunoreactive cells in the SFO. These results, beyond demonstrating once again the effectiveness of this perinatal programming paradigm, allow us to suggest that depending on which systems are mainly stimulated or inhibited during perinatal programming, particular neural pathways and sensors are modified during early ontogeny. Therefore, according to the specific characteristics of the perinatal environment, adults drank larger or smaller volumes of fluids than their controls when they were exposed to a specific challenge that recruits essential sensors and circuits.

Another perinatal programming model, which induces maternal natriophilia but in conjunction with polydipsia, is the partial aortic ligation (PAL). This procedure induces a decrease in the blood flow of the ischemic kidney sensed by the renal volume receptors, signaling the local hypovolemia that finally leads to a chronic secretion and release of renin, over-activating the renin-angiotensin system (RAS). In this sense, PAL is an experimental paradigm characterized by polydipsia, natriophylia, arterial blood pressure (PSA), and urinary volume increases (Costales et al., 1984). PAL-dams exhibited an increment in sodium and water intake and in plasma renin activity levels compared to pregnant females with access to hypertonic sodium but with sham ligation (Argüelles et al., 2000; Perillan et al., 2004, 2007).

Several effects have been reported in PAL-offspring, such as: i) changes in aldosterone levels and by inference on others RAS components (Vijande et al., 1996); ii) increases in basal and isoproterenol-induced water intake and iii) elevated levels of hypotonic sodium intake (Argüelles et al., 2000; Perillan et al., 2004). However, it should be noted that these antecedents analyze the PAL programming effects in comparison to sham-ligation but both groups are exposed to hypertonic sodium solution. Nevertheless, such as aforementioned: free access to hypertonic sodium is able to program the offspring by itself. However, to our knowledge, there are no comparative studies that discriminate which are the fluid intakes changes in adult offspring of dams with availability of hypertonic sodium chloride solution and which of PAL dams.

The main goals were i) to characterize fluid intake, plasma electrolyte, PRA levels, and kidney atrophy in dams and/or female pups as a function of the perinatal manipulation (PM) models; ii) to compare drinking responses induced by three different models of thirst in adult programmed-offspring. The thirst models studied in adult offspring were absolute dehydration induced by overnight water deprivation, relative dehydration induced by a hypertonic intravenous infusion of NaCl and hypovolemic thirst induced by sodium depletion. We analyzed the fluid intake and excretion patterns as a function of the PM paradigms. Also, in the hypovolemic thirst model, we examined the brain cell activity patterns in nuclei of the *lamina terminalis* (LT), hypothalamus, and brainstem. We postulate that the perinatal programming is a phenomenon that differentially affects particular systems and target sites inducing specific drinking and brain cell activity responses depending on the particular environmental conditions present during the perinatal programming.

## 2. Materials and Methods

### 2.1 Animals

Wistar-derived rats born and reared at the vivarium of the Ferreyra Institute (INIMEC-CONICET-UNC, Argentina) were employed. The animal colony was kept at 22-24 °C under artificial lighting conditions (lights on: 08:00–20:00 h). Female rats weighing 220–250g, 70–75 days old and non-littermates were individually housed in standard holding chambers (40×40×70cm). The Principles of Laboratory Animal Care were followed and animals used in this study were maintained and treated according to the Guidelines for the Care and Use of Mammals in Neuroscience and Behavioral Research (National Research Council 2003) and also to the Institutional Committee of Laboratory Animal Use and Care of our institution (CICUAL-INIMEC-CONICET-UNC).

Female rats were randomly assigned to receive one of the corresponding perinatal manipulation (PM) models. A PM experimental group [**PAL-W/Na group]** received a partial aortic ligation (PAL) and free access to tap water, hypertonic sodium chloride solution (0.45 M NaCl) and standard commercial diet (Cargill Inc. Argentina, containing approx. 0.18% NaCl). Another PM experimental group **[Sham-W/Na group]** received a sham-PAL ligation and free access to tap water, hypertonic sodium chloride solution, and commercial diet. As a PM control group, females received a sham-PAL ligation in addition to free access to tap water only and commercial diet **[Sham-W group]**. Exposure periods to hypertonic sodium chloride solution and other details of the experimental design were described in **Figure 1**. Briefly, after a week of adaptation to the individual holding chamber and the hypertonic sodium solution, the corresponding sham-PAL or PAL surgery was performed in females. After one week of surgery recovery, females were placed with males for mating and once the pregnancy was confirmed, females were returned to the individual home-cage. Females and their litters were individually kept with the corresponding solutions (water and/or hypertonic sodium solution) throughout pregnancy and lactation. Within 24 h after birth, litters were culled to 10 pups (5 males and 5 females whenever possible). Litters with less than six pups were not included. At weaning (postnatal day PD 21–22), plasma from both dams and their female pups, and maternal kidneys were collected to further analysis. Only the male pups continued the experiments receiving the same conditions as their dam until they reach one month of life (PD 28). From then on, males of both experimental conditions were kept under standard conditions of water and food until 2 months of age (PD 60–70), such as PM control group.

**Figure 1:**
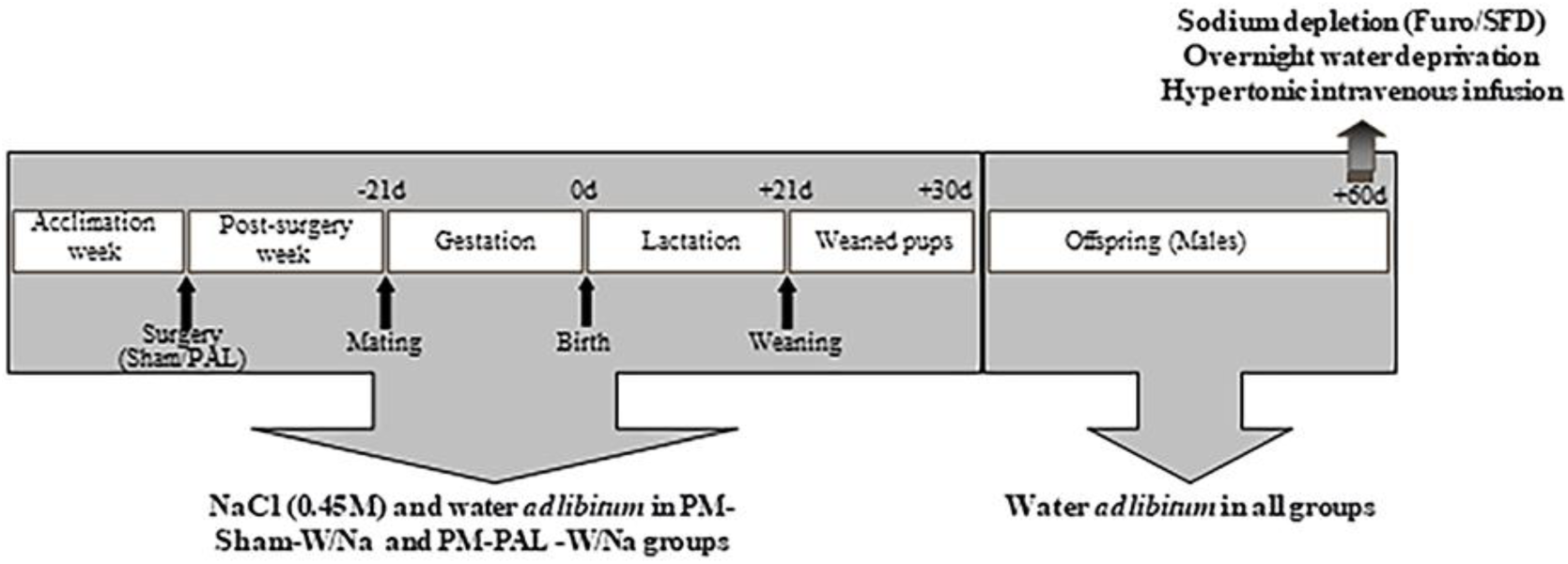
Schematic diagram representing the experimental design employed to the perinatal manipulation (PM) models. Note that from PD28 until PD60, males of each PM group were kept on standard conditions.

### 2.2 Experiment #1: Characterization of the PM models in dams and female pups: Effects of PAL and the maternal exposure to hypertonic sodium chloride solution during gestation and lactation

#### 2.2.1 Experiment #1.a

Maternal water and hypertonic sodium chloride solution intake With the aim to characterize the drinking patterns during the pregnancy and lactation of dams of each PM models, water, hypertonic sodium solution (0.45 M) and total fluid intakes were recorded from the week of adaptation and throughout pregnancy and lactation. The maternal intake of water and hypertonic sodium solution of twenty-two dams (Sham-W = 9; Sham-W/Na = 6 and PAL-W/Na = 7) were daily recorded and weekly averaged, except to the complete pregnancy period in which each intake was averaged taking into account the total duration of pregnancy (∼20-22 days). Data are shown as average of fluid consumed/dam (water, sodium, and total fluid intake) during the following periods: adaptation week; first week of pregnancy; complete pregnancy period; first, second, and third lactation weeks. Dams’ intake during mating days was not taken into account to calculate the volume drunk of any period. The duration of the pregnancy (gestational days) and the litter size on the day of delivery were also recorded.

#### 2.2.2 Experiment #1.b

Plasma electrolyte concentration, osmolality, and protein assays in dams and female pups at weaning In order to determine the PM effects on plasma electrolyte concentration ([Na+], [K+] and [Cl-]), osmolality, and volemia at weaning, trunk blood of dams and female pups was collected by decapitation in plastic tubes containing EDTA (final concentration 2mg/mL blood) and immediately centrifuged at 4°C for 20min at 3,000g. Then, plasma was removed and kept at -20°C until determination (note that pooled blood was not necessary for the measurements in pups). Plasma sodium [Na+], potassium [K+], and chloride [Cl-] concentrations were determined using an Ion Selective Electrode (Hitachi Modular P+ISE. Roche 8 Diagnostic). Plasma osmolality was analyzed by vapor pressure osmometry (VAPRO 5520) and volemia was indirectly inferred by the plasma protein concentration which was measured in an absorbance microplate reader (BioTek EL800), according to the protocol proposed by Lowry et al. 1951.

#### 2.2.3 Experiment #1.c

Reduction of the ischemic kidney and plasma renin activity (PRA) in PAL-W/Na dams at weaning With the purpose of quantify the partial aortic ligation (PAL) effects in ligated-dams, we assessed the reduction of the ischemic kidney (kidney subjects to ligation) as well as the level of compensatory expansion in the opposite kidney. At weaning, we removed and weighed both maternal kidneys and calculated the percentage of reduction of one of them compared to the other. To standardize the ligation procedure, only PAL dams with a percentage of reduction between 40% and 60% relative to each other were included in the experiments.

Furthermore, maternal PRA levels were quantified. At weaning, maternal plasma samples were collected as previously described (Experiment 1.b) and kept at - 20°C until determination. PRA was measured by radioimmunoassay (RIA) of angiotensin I (DiaSorin, Saluggia, Italy) in the presence of reagents that inhibit angiotensin I-converting enzyme and angiotensinases. Intra- and inter-assay coefficients of variation were lower to 8% and 6.66%, respectively.

### 2.3 Experiment #2: Long-term PM effects on adult offspring challenged with an absolute dehydration induced by overnight water deprivation

Thirty-two adult males (≥PD60) were placed in individual metabolic cages with access to normal food and deionized water in graduated tubes for ∼30 hours (acclimation period). On the next day, water was removed (∼ 5.00 PM) to achieve water deprivation, leaving animals with an ad libitum access to food for the following ∼17 hours (deprivation period). The next morning, cumulative volumes of water ingested were measured at 60-min after a water intake test. To measure osmolality and urinary concentration of electrolytes ([Na+], [K+], [Cl-]), urines were collected during each period: acclimation, deprivation, and test. Urinary volumes were adjusted to 100g of body weight and the corresponding collection time (30 h to acclimation period, 17 h to water deprivation period, and 2 h to test period).

### 2.4 Experiment #3: Long-term effects of the PM in adult offspring in response to a relative dehydration induced by a hypertonic intravenous infusion

Sixty adult males (≥ PD 60) were anesthetized and subjected to a femoral vein cannulation. Then, animals were placed in individual metabolic cages with ad libitum access to food and deionized water. With the aim of controlling the basal status of the animals before the intake test, body weight on the day of surgery (BW-1) and the day of infusion (BW-2), as well as the overnight water intake on the night before the iv infusion day, were recorded. After 5-10 minutes of acclimation, animals were infused with a hypertonic (1.5M NaCl) or isotonic (0.15M NaCl) solution (0.15mL/min iv for 20 minutes) with an infusion pump (SyringePump, NE1800) and according to the protocol of Ho et al. 2007. Cumulative water intake was recorded 100 minutes after the beginning of the infusion (20 min corresponding to infusion time and 60 min post-infusion). Urine collection was performed 140 minutes after the start of the infusion.

### 2.5 Experiment #4: Long-term PM effects in adult offspring challenged by a hypovolemic thirst model

To assess adult fluid intake induced by a hypovolemic thirst model as a function of the PM models, seventy-four adult males (≥PD60) were placed in individual metabolic cages with access to commercial diet, deionized water, and hypertonic sodium solution (0.45 M NaCl) in graduated tubes. After a 48-hour acclimation period of spontaneous intake of both solutions, males of each PM group were randomly assigned to the sodium-replete **[Veh]** or Furosemide–sodium depleted **[Furo]** group. Sodium depletion was carried out by a combined treatment of Furosemide and sodium-free diet (SFD) according to the protocol of Sakai, Nicolaïdis, & Epstein, 1986. Furosemide was administered to the animals (Lasix, Sanofi Aventis lab, sc; 40mg/kg) in 2 injections separated by 2 h (at noon). At the time of the first injection, commercial diet was replaced by SFD, and hypertonic sodium chloride solution was withdrawn. Sodium-replete animals were injected with the equivalent volume of vehicle solution (0.15 M NaCl) and had access to the commercial diet. Twenty hours after, animals were subjected to hypertonic sodium solution and water intake test of 60 minutes of duration, during which time the cumulative ingested volumes of both solutions were measured.

### 2.6 Experiment #5: Long-term PM effects on brain activation patterns (Fos + cells) in adult offspring exposed to a hypovolemic thirst model

A total of fourteen males (≥PD60, n ≥ 3/experimental group) were administered with 40 mg/kg of Furosemide, via sc. As previously described, males were injected with Furosemide (Lasix, Sanofi Aventis lab, sc) in two injections separated by 2 h (at noon). At the time of the first injection, commercial diet was replaced by SFD and hypertonic sodium chloride solution was withdrawn. Twenty hours after sodium depletion, animals were perfused without access to the intake test.

### 2.7 Partial aortic ligation

Adult females were anesthetized with ketamine-xylazine (1.25mL/kg, sc) and subjected to PAL or sham-PAL surgery, as described Costales et al. 1984. First, the abdominal area was shaved and a longitudinal incision was made in the midline. The abdominal aorta was dissected between the renal arteries. At an intermediate point between two renal arteries - below the mesenteric artery and above the left renal artery - a metal stylet of 0.5 mm of diameter was placed next to the aorta and this aggregate was ligated with a 4/0 suture thread. The stylet was then removed and the incision was closed in layers.

### 2.8 Femoral vein cannulation

Adult males were anaesthetized with ketamine–xylazine (1.25 mL/kg, sc) for cannulation surgery, as described Caeiro and Vivas 2008. The right leg area was shaved and a horizontal incision was made. A polyethylene PE-10 catheter (0.02500 OD-0.01100 ID, Clay Adams, Parsippany, New Jersey, Jersey, USA) with heparin–saline (2 IU/mL) was implanted into the left femoral vein welded to a PE-50 (0.03900 OD-0.02300 ID, Clay Adams, Parsippany, New Jersey, USA). The catheter was subcutaneously conducted to an exit between the shoulder blades and it was connected to a modified needle, in order to prevent movement of the catheter, and then it was sealed.

### 2.9 Immunohistochemistry

Animals were anesthetized with chloral hydrate 6% (0.6mL/100g bw, ip) and transcardially perfused with ∼100 mL of normal saline solution followed by ∼400 mL of 4% paraformaldehyde in 0.1 M phosphate buffer (PB, pH 7.2). Brains were removed, fixed overnight in the perfusion solution and stored at 4 °C in PB containing 30% sucrose. Free-floating 40 μm coronal sections were cut using a freezing microtome. Immediately before the immunostaining, the sections were placed in a solution with 10% H_2_O_2_ and 10% methanol for 60 min. Then, an incubation was performed in a 10% normal horse serum (NHS-Gibco, Auckland, NZ) for 1 h in PB to block non-specific binding sites. Fos immunoreactivity (Fos-ir) was detected using a standard avidin–biotin peroxidase protocol. Free-floating sections were incubated overnight at room temperature in an antibody raised in rabbit against a synthetic 14 amino acid sequence corresponding to residues 4–17 of human Fos (Ab-5, Oncogene Science, Manhasset, NY) diluted 1:10,000 in a PB solution containing 2% NHS and 0.3% Triton X-100 (Sigma Chemical Co., St. Louis, MO, USA). After washing in PB, the sections were incubated in biotin-labeled anti-rabbit immunoglobulin (Jackson Immunoresearch Laboratories) diluted 1:500 in 1% NHS-PB and in a avidin–biotin peroxidase complex (Vector Laboratories Inc., Burlingame, CA, USA) diluted 1:200 in 1% NHS-PB for 1 h at room temperature. The peroxidase label was detected using diaminobenzidine hydrochloride (DAB; Sigma Chemical Co., St. Louis, MO, USA) intensified with 1% cobalt chloride and 1% nickel ammonium sulfate. This method produces a blue-black nuclear reaction product. Free-floating sections were mounted on gelatinized slides (with Albrecht’s gelatine), air-dried overnight, dehydrated, cleared in xylene, and placed under a coverslip with DPX mounting medium (Fluka, Buchs, Switzerland).

### 2.10 Cytoarchitectural and quantitative analysis

Brain nuclei were identified and delimited on the basis of the plates from the rat brain atlas of Paxinos and Watson 2007. The number of Fos-ir + cells was counted at the distance from bregma as it is indicated between brackets: *organum vasculosum of the lamina terminalis* (OVLT, -0.20 mm), median preoptic nucleus (MnPO, -0.40 mm), subfornical organ (SFO, -0.92 mm), supraoptic nucleus (SON, -1.3 mm), ventrodorsal and ventrolateral subdivisions of the dorsal raphe nucleus (DRN/VD and DRN/VL, - 8.00 mm), lateral parabrachial nucleus (LPB) and locus coeruleus (LC, -9.3 mm), area postrema (AP), and nucleus of the solitary tract (NTS, -13.68 mm). Different subnuclei of paraventricular nucleus (PVN) were quantified as follow: medial magnocellular (PaMM, -1.40 mm); lateral magnocellular (PaLM) and ventral (PaV) subdivisions (−1.80 mm); and posterior paraventricular subnucleus (PaPo), and medial parvocellular (PaPM, -2.12 mm). Fos-ir quantification was performed using a computerized system that included a Zeiss microscope equipped with a DC 200 Leica digital video camera attached to a contrast enhancement device. Video images were digitized and analyzed using Adobe Photoshop Image Analysis Program CS2, version 9. Representative sections were acquired under the same illumination conditions. The counting was carried out in three to five animals of each experimental condition. The investigator who conducted the counting of Fos-ir + cells was blinded to the experimental groups. Because one section of each nucleus was quantified, no corrections were necessary to avoid double counting.

### 2.11 Statistical analysis

All data are expressed as mean +/- SE. All variables were analyzed by analysis of variance (ANOVA) with repeated measures (period or solution) when appropriate. Both main effects and interactions were considered to be statistically significant at p < 0.05. Partial eta-squared (η2p) was used to describe effect sizes of ANOVAs, which were interpreted using the following guidelines [small (η2p = 0.01–0.05), medium (η2p = 0.06 – 0.13), and large (η2p = ≥ 0.14) (Lakens, 2013)]. Post hoc comparisons were performed using Least Significant Difference (LSD) tests. Linear correlations were calculated to assess the relationship between total water consumed and excreted urine volumes in the test of the absolute dehydration protocol. Differences between specific Pearson’s correlation coefficients were statistically analyzed using z tests. Analyses were performed utilizing Statistica Stat-Soft Inc. version 8.0 Tulsa, OK, USA.

## 3. RESULTS

### 3.1 Experiment #1: Characterization of the PM models in dams and female pups: Effects of PAL and maternal exposure to hypertonic sodium chloride solution during gestation and lactation

#### 3.1.1 Experiment #1.a

Maternal fluid intake (water, hypertonic sodium solution-0.45 M and total fluid)

Maternal water, hypertonic sodium solution, and total fluid intake were analyzed across the following periods: adaptation week [Adapt], first week of pregnancy [1^st^ Pregn], complete pregnancy [Compl Pregn], and during the 3 weeks of lactation [1^st^, 2^nd^, and 3^rd^ Lact]. Analyses were performed using a 2-way ANOVA with repeated measures between periods.

##### Maternal water intake

As shown in **Table 1**, the ANOVA indicated significant main effects of PM and period factors [F (2, 19) = 4.20, p = 0.0309, η2p = 0.31; F (5, 95) = 64.93, p < 0.0001, η2p = 0.77, respectively]. However, a non-significant interaction between the factors was observed. Post hoc comparisons showed that PAL-W/Na dams consumed greater water volumes than Sham-W and Sham-W/Na dams, independently to the periods (Sham-W = 97.89+/- 8.53; Sh-W/Na = 86.05+/-10.45; PAL-W/Na = 125.53+/-9.68). Also, post hoc comparisons indicated that, beyond to PM model analyzed, maternal water intake of each period was higher than the previous period, except between 1^st^ Pregn and Compl Pregn periods.

**Table 1.** Maternal water, sodium hypertonic and total fluid intake [mL/day/dam] across periods that involve the PM protocols.

| *Maternal intake* |  |  | Average of volume consumed (mL/day/dam) |  |  |
| --- | --- | --- | --- | --- | --- |
|  |  |  | Sham-W | Sham-W/Na | PAL-W/Na |
| *Water* | *Period* | **Adapt** | 40.0 +/- 5.2 (9) | 31.9 +/- 2.9 (6) | 41.1 +/- 3.5 (7) |
|  |  | **1<sup>st</sup> Pregn** | 55.8 +/- 5.3 (9) | 61.4 +/- 9.9 (6) | 76.3 +/- 7.3 (7) |
|  |  | **Compl Pregn** | 72.0 +/- 5.4 (9) | 70.1 +/- 7.6 (6) | 95.9 +/- 11.2 (7) |
|  |  | **1<sup>st</sup> Lact** | 97.8 +/- 7.9 (9) | 84.9 +/- 21.6 (6) | 124.7 +/- 13.9 (7) |
|  |  | **2<sup>nd</sup> Lact** | 134.0 +/- 12.7 (9) | 120.5 +/- 27.1 (6) | 188.8 +/- 28.9 (7) |
|  |  | **3<sup>rd</sup> Lact** | 187.8 +/- 23.0 (9) | 147.6 +/- 24.2 (6) | 226.4 +/- 26.4 (7) |
| *Sodium hypertonic* | *Period* | **Adapt** | NA | 10.1 +/- 1.3 (6) | 6.5 +/- 1.2 (7) |
|  |  | **1<sup>st</sup> Pregn** | NA | 22.7 +/- 4.5 (6) | 21.2 +/- 4.2 (7) |
|  |  | **Compl Pregn** | NA | 20.7 +/- 3.7 (6) | 29.3 +/- 3.5 (7) |
|  |  | **1<sup>st</sup> Lact** | NA | 10.6 +/- 4.0 (6) | 27.1 +/- 3.7 (7) |
|  |  | **2<sup>nd</sup> Lact** | NA | 18.5 +/- 5.9 (6) | 25.9 +/- 5.5 (7) |
|  |  | **3<sup>rd</sup> Lact** | NA | 18.4 +/- 3.9 (6) | 24.4 +/- 3.6 (7) |
| *Total fluid* | *Period* | **Adapt** | 40.0 +/- 5.2 (9) | 43.7 +/- 3.4 (6) | 47.5 +/- 3.4 (7) |
|  |  | **1<sup>st</sup> Pregn** | 55.8 +/- 5.3 (9) | 92.8 +/- 19.9 (6) | 97.5 +/- 8.2 (7) |
|  |  | **Compl Pregn** | 72.0 +/- 5.4 (9) | 100.8 +/- 14.3 (6) | 125.2 +/- 12.6 (7) ++ |
|  |  | **1<sup>st</sup> Lact** | 97.8 +/- 7.9 (9) | 95.5 +/- 22.7 (6) | 151.7 +/- 12.6 (7) ** |
|  |  | **2<sup>nd</sup> Lact** | 134.0 +/- 12.7 (9) | 139.0 +/- 27.2 (6) | 214.7 +/- 34.3 (7) ** |
|  |  | **3<sup>rd</sup> Lact** | 187.8 +/- 23.0 (9) | 166.0 +/- 26.5 (6) | 250.7 +/- 29.0 (7) ** |

Values are expressed as means +/- SE, (n). Periods: adaptation week [*Adapt*], first week of pregnancy [*1<sup>st</sup> Pregn*], complete pregnancy [*Compl Pregn*] and lactation weeks [*1<sup>st</sup>*, *2<sup>nd</sup>*, and *3<sup>rd</sup> Lact*]. (++) Significant differences between PAL-W/Na with respect to Sham-W group, $p < 0.01$ . (\*\*) Significant differences between PAL-W/Na with respect to the other groups, $p < 0.01$ . NA, not applicable.

##### Maternal sodium hypertonic (0.45M NaCl) intake

The ANOVA indicated a significant main effect of period [F (5, 55) = 5.76, p = 0.0002, η2p = 0.34]. No significant effect of PM factor or interaction between factors were observed. However, despite the fact that the interaction between PM and period did not reach statistically significant levels [F (5, 55) = 2.22, p = 0.0655], a high hypertonic sodium intake was observed in PAL-W/Na dams during periods that included the complete pregnancy and also the 1^st^ and 2^nd^ weeks of lactation. An increase of 41.77% in the complete pregnancy, 155.52% in 1^st^ week of lactation and 40.03% in the 2^nd^ week of lactation was observed in PAL-W/Na dams compared to Sham-W/Na dams (**Table 1)**. Regarding the hypertonic sodium intake across the periods, post hoc comparisons revealed significant increments between the adaptation week relative to the other periods.

##### Maternal total fluid intake

Main effects of PM and period were observed [F (2, 19) = 6.14, p = 0.0088, η2p = 0.39, and F (5, 95) = 61.93, p < 0.0001, η2p = 0.77, respectively], as well as a significant interaction between them [F (10, 95) = 2.36, p = 0.0152, η2p = 0.20]. Post hoc comparisons indicated a gradual increase in the total intake across the periods. However, focusing on the analysis of the total fluid intake of different PM dams within each period, it is observed that PAL-W/Na dams consumed high volumes of fluid relative to Sham-W dams when considering the complete pregnancy period. This group of dams also continued drinking higher fluid volumes during each lactation week relative to both Sham-W and Sham-W/Na dams (**Table 1**).

Regarding the pregnancy duration and litter sizes recorded on delivery day, no significant differences were found between the PM models. The means +/- SE for days of pregnancy were Sham-W = 22.10 +/- 0.18; Sham-W/Na = 22.20 +/- 0.36 and PAL-W/Na = 22.30 +/- 0.33. Litter sizes were Sham-W = 10.33 +/- 1.02; Sham-W/Na = 9.83 +/- 0.75 and PAL-W/Na = 8.33 +/- 0.67.

#### 3.1.2 Experiment #1.b

Plasma electrolyte concentration, osmolality, and protein assays in dams and female pups at weaning One-way ANOVAs were conducted to analyze plasma electrolyte concentration, osmolality, and protein concentration.

##### Dams at weaning

A main effect of the PM factor was observed by K+ and osmolality parameters [F (2, 11) = 6.18, p = 0.0159, η2p = 0.53, and F (2, 12) = 5.68, p = 0.0184, η2p = 0.49, respectively]. Post hoc comparisons showed a reduction in plasma K+ concentration in PAL-W/Na dams in relation to Sham-W/Na dams and also a decrease in plasma osmolality in PAL-W/Na dams with respect to another both PM conditions **(Table 2)**. No significant differences in plasma levels of Na+, Cl-, and plasma protein concentration were observed in dams of any PM groups.

**Table 2.**
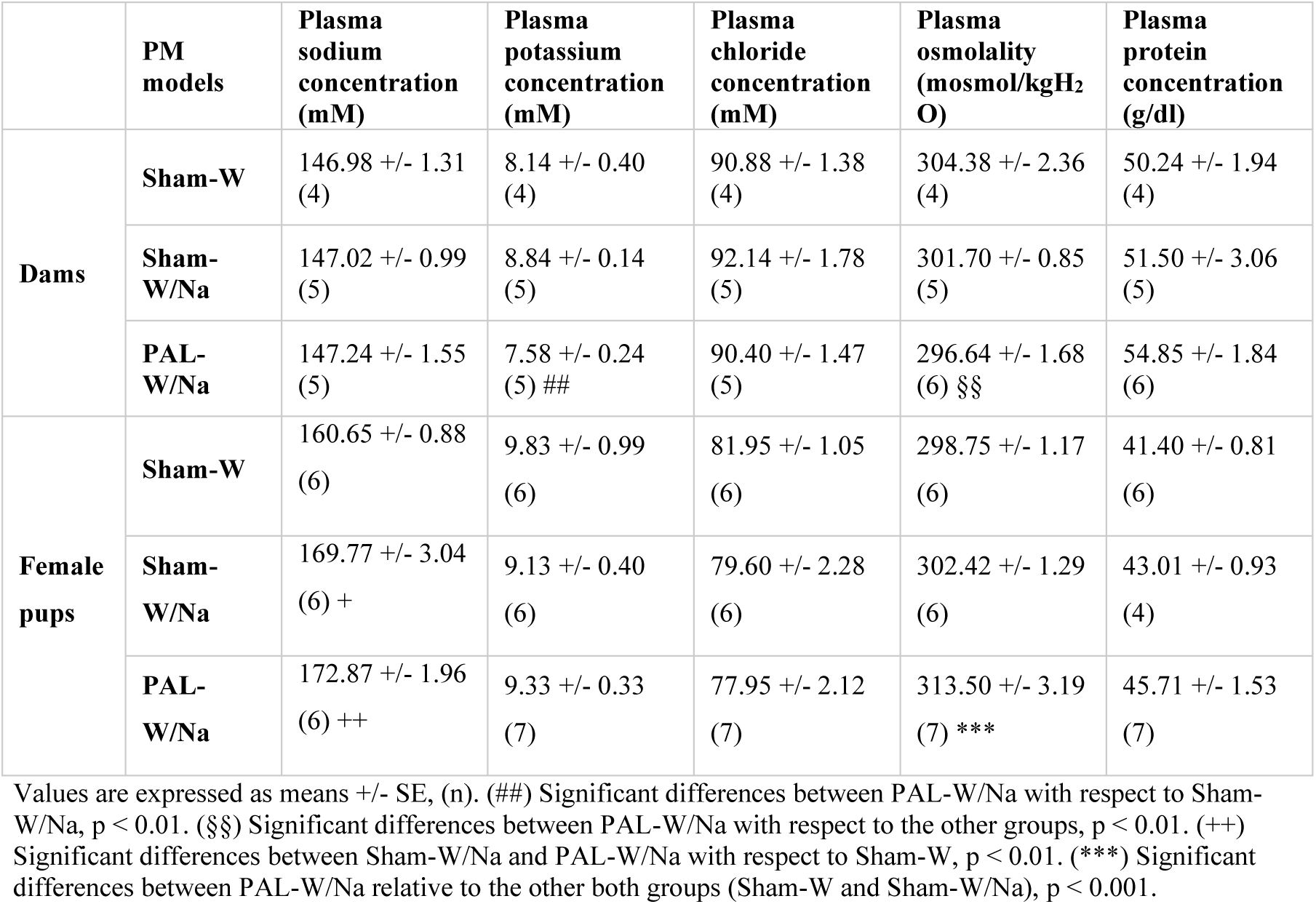
Plasma electrolyte concentration (Na+, K+, and Cl-), osmolality, and protein concentration in dams and female pups at weaning (PD 21) as a function of the PM models.

|  | PM models | Plasma sodium concentration (mM) | Plasma potassium concentration (mM) | Plasma chloride concentration (mM) | Plasma osmolality (mosmol/kgH<sub>2</sub>O) | Plasma protein concentration (g/dl) |
| --- | --- | --- | --- | --- | --- | --- |
| Dams | Sham-W | 146.98 ± 1.31 (4) | 8.14 ± 0.40 (4) | 90.88 ± 1.38 (4) | 304.38 ± 2.36 (4) | 50.24 ± 1.94 (4) |
| | Sham-W/Na | 147.02 ± 0.99 (5) | 8.84 ± 0.14 (5) | 92.14 ± 1.78 (5) | 301.70 ± 0.85 (5) | 51.50 ± 3.06 (5) |
| | PAL-W/Na | 147.24 ± 1.55 (5) | 7.58 ± 0.24 (5) ## | 90.40 ± 1.47 (5) | 296.64 ± 1.68 (6) §§ | 54.85 ± 1.84 (6) |
| Female pups | Sham-W | 160.65 ± 0.88 (6) | 9.83 ± 0.99 (6) | 81.95 ± 1.05 (6) | 298.75 ± 1.17 (6) | 41.40 ± 0.81 (6) |
| | Sham-W/Na | 169.77 ± 3.04 (6) + | 9.13 ± 0.40 (6) | 79.60 ± 2.28 (6) | 302.42 ± 1.29 (6) | 43.01 ± 0.93 (4) |
| | PAL-W/Na | 172.87 ± 1.96 (6) ++ | 9.33 ± 0.33 (7) | 77.95 ± 2.12 (7) | 313.50 ± 3.19 (7) *** | 45.71 ± 1.53 (7) |

Values are expressed as means ± SE, (n). (##) Significant differences between PAL-W/Na with respect to Sham-W/Na, $p < 0.01$ . (§§) Significant differences between PAL-W/Na with respect to the other groups, $p < 0.01$ . (++) Significant differences between Sham-W/Na and PAL-W/Na with respect to Sham-W, $p < 0.01$ . (\*\*\*) Significant differences between PAL-W/Na relative to the other both groups (Sham-W and Sham-W/Na), $p < 0.001$ .

##### Female pups

A significant effect of the main factor PM was observed by Na+ and osmolality pup’s parameters [F (2, 15) = 8.71, p = 0.0031, η2p = 0.54, and F (2, 16) = 11.99, p = 0.0006, η2p = 0.60, respectively]. A posteriori comparisons showed an increase in plasma Na+ concentration in female pups subjected to perinatal exposure to a hypertonic sodium solution, regardless of the presence or absence of ligation (PAL-W/Na and Sham-W/Na). Besides, post hoc comparisons revealed an increase in plasma osmolality only in PAL-W/Na pups. No significant differences were observed in the plasma K+ and Cl-levels in pups. In the case of plasma protein concentration, a tendency was found in PAL-W/Na female pups [F (2, 16) = 3.46, p =0.0564, η2p = 0.30] where protein levels were high in PAL-W/Na female pups relative to the other PM groups **(Table 2)**.

#### 3.1.3 Experiment #1.c

Reduction of the ischemic kidney and plasma renin activity (PRA) in PAL-W/Na dams at weaning

Both kidneys of the dams of all PM models were weighed at weaning day. A 2-way ANOVA with repeated measures was performed. A significant effect of both factors: PM and weight of the kidneys [F (2, 49) = 6.11, p = 0.0043, η2p = 0.20, and F (1, 49) = 133.64, p < 0.0001, η2p = 0.73, respectively] as well as a significant interaction between them [F (2, 49) = 122.31, p < 0.0001, η2p = 0.83] were observed. A posteriori comparisons indicated that both right and left kidneys of PAL-W/Na mothers weighed different from their counterpart in Sham-W/Na and Sham-W dams. Specifically, PAL-W/Na dams showed a significant hypertrophy of the right kidney and a strong atrophy of the left kidney (**Figure 2A**). In addition, when analyzing the percentage of reduction of one kidney with respect to the another, the 1-way ANOVA showed, as expected, that the PAL effectiveness (calculated through the percentage of renal atrophy in PAL-W/Na mothers) was between 40 and 60% [F (2, 49) = 165.22, p < 0.0001, η2p = 0.87, (**Figure 2A**)].

**Figure 2:**
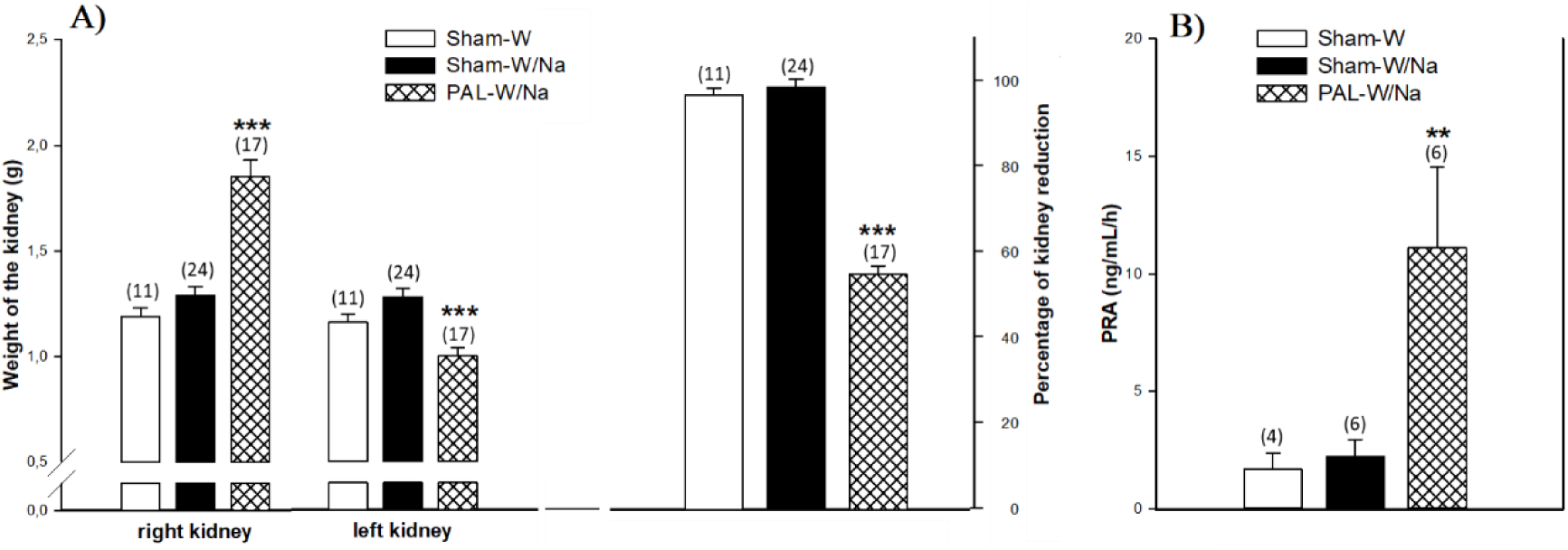
**A)** Kidney weights and percentage of kidney reduction in dams at weaning as a function of PM conditions. **B)** Plasma renin activity (PRA) in dams at weaning as a function of PM conditions. Values are expressed as means +/- SE, (n). Significant differences between PAL-W/Na relative to the other both groups (Sham-W and Sham-W/Na), (^***^) p <0.001 and (^**^) p < 0.01, respectively.

PRA levels were determined in maternal blood samples obtained at the weaning day and they were analyzed via 1-way ANOVA. A significant effect of the PM factor was observed [F (2, 13) = 5.43; p < 0.0193, η2p = 0.46]. As shown in **Figure 2B**, a posteriori comparisons indicated a significant increase in PRA levels in PAL-W/Na mothers compared to the other 2 groups (Sham-W/Na and Sham-W).

### 3.2 Experiment #2: Long-term PM effects on adult offspring challenged with an absolute dehydration induced by overnight water deprivation

#### Water intake test

Water intake test after an overnight water deprivation as a function of PM models was analyzed via 1-way ANOVA. Significant differences were found in water volumes between both PM models (Sham-W/Na and PAL-W/Na) with respect to the control PM group (Sham-W) [main effect F (2, 29) = 5.55, p = 0. 0091, η2p = 0.28]. Post hoc comparison indicated that Sham-W/Na and PAL-W/Na offspring consumed lower volumes of water than Sham-W group (**Figure 3A**).

**Figure 3:**
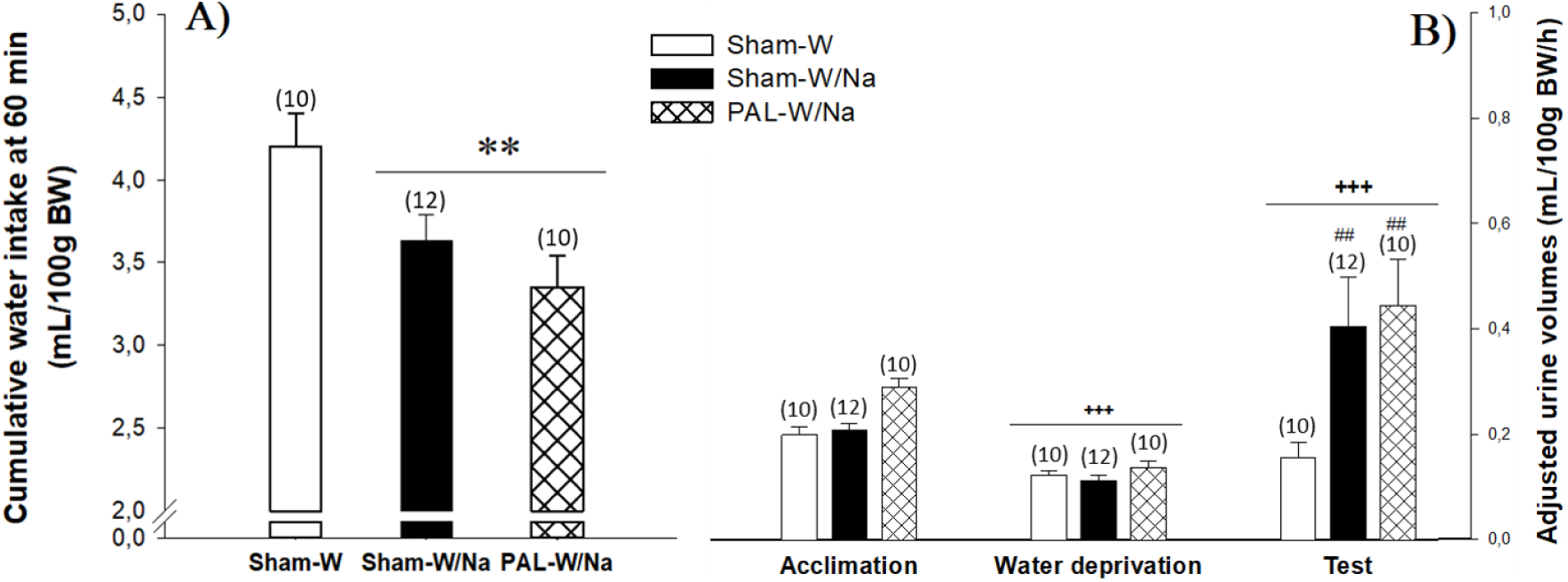
**A)** Cumulative water intake at 60 min (mL/100g BW) after an overnight water deprivation in adult offspring of each PM model. **B)** Adjusted urine volumes (mL/100g BW/hour) excreted at each collection period: acclimation, water deprivation and intake test. Values are expressed as means +/- SE, (n). (^**^) Significant differences between Sham-W/Na and PAL-W/Na groups with respect to Sham-W group, p < 0.01. (+++) Significant differences between collection periods, p < 0.0001. (##) Significant differences between Sham-W/Na and PAL-W/Na groups with respect to Sham-W group, p < 0.01.

#### Excreted urine volumes

Adjusted urine volumes per hour (mL/100g BW/h) at each collection period were analyzed via 2-way ANOVA with repeated measure (collection period). PM and collection period achieved significance [F (2, 29) = 5.47, p = 0.0096, η2p = 0.27, and F (2, 58) = 15.70, p < 0.0001, η2p = 0.35, respectively]. LSD test showed a significant reduction in excreted urine volume/h during the overnight water deprivation period and a significant increment in this parameter during the intake test period (**Figure 3B**). A 2-way interaction between PM and collection period also achieved significance [F (4, 58) = 3.22, p = 0.0186, η2p = 0.18]. A posteriori comparisons indicated that Sham-W/Na and PAL-W/Na groups excreted higher urine volumes by hour than Sham-W animals (**Figure 3B**). Taking into account that these two PM groups also showed a reduced water intake during the test, it is possible to postulate that the hydric status of these animals has not normally been re-established.

In order to analyze the relationship between the total water consumed and excreted urine volumes in the test as a function of each PM model, linear correlations were calculated (Pearson correlation coefficient). A significant positive correlation was found between these two variables in PAL-W/Na’s adult offspring [r = .70, n = 10, p = .0235] (**Figure 3C**). A similar profile of water consumed/excreted urine was observed in the case of Sham-W/Na animals, although it did not reach statistically significant levels [r = .54, n = 12, p = .0725]. These analyzes allowed to observe that, while the control animals retained everything they drank (x-axis variation - mL of water consumed-did not modify y-axis values - mL of urine excreted), the descendants of both PM models had a linear correlation between drinking/excretion, suggesting a lower water retention.

**Figure 3:**
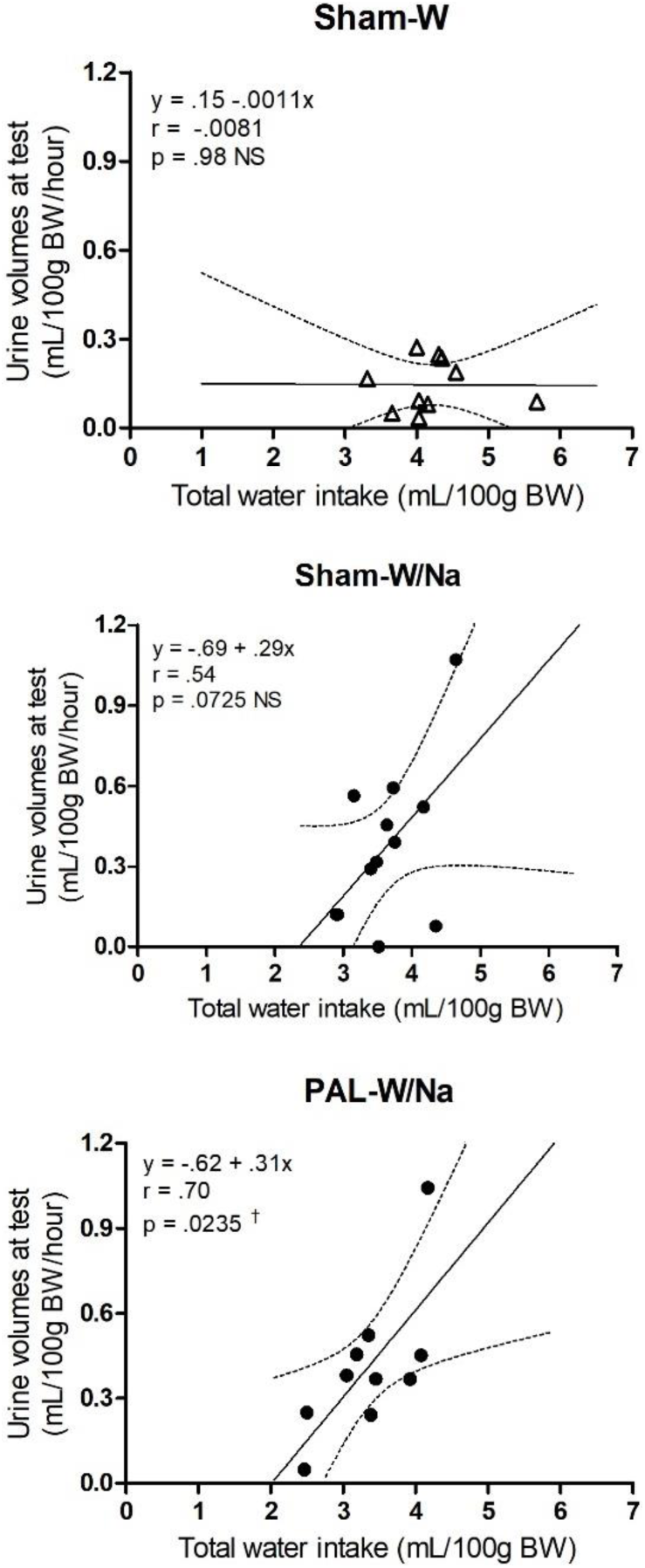
**C)** Lineal correlation between total water intake at 60 min (mL/100g BW) and adjusted urine volumes in the test (mL/100g BW/hour) as a function of Sham-W (top panel), Sham-W/Na (middle panel) and PAL-W/Na (bottom panel) groups. (†) Significant correlations between variables under consideration in the PAL-W/Na group, p < 0.05.

### 3.3 Experiment #3: Long-term effects of the PM in adult offspring in response to a relative dehydration induced by a hypertonic intravenous infusion

Sixty-min water intake test after the intravenous infusion as a function of PM model and solution infused were analyzed via 2-way ANOVA. A significant increase in water intake was observed in animals infused with a hypertonic NaCl (HP) solution with respect to those infused with an isotonic NaCl (ISO) solution [Solution infused-main effect: F (1, 54) = 45.43, p < 0.0001, η2p = 0.46]. In relation to the PM effects, a significant main effect of PM model **(inset in Figure 4A)** and a significant interaction between this factor and solution infused were found [F (2, 54) = 3.34, p = 0. 0428, η2p = 0.11, and F (2, 54) = 4.15, p = 0.0211, η2p = 0.13, respectively]. A posteriori comparisons indicated a significant increase in water intake induced by the hypertonic NaCl infusion in Sham-W/Na group compared to Sham-W and PAL-W/Na groups (**Figure 4A**). Non-significant differences were found between animals of the different PM models when were infused with isotonic solution, in terms of body weight at the day of the surgery (BW-1) and at the day of infusion (BW-2) as well as the overnight water intake **(Table 3)**.

**Table 3.** Body weight at the day of the surgery (BW-1) and at the day of the infusion (BW-2), and the overnight water intake before the infusion in adult male offspring.

| PM conditions | BW-1 (g) | BW-2 (g) | Overnight water intake (mL/100 g BW) |
| --- | --- | --- | --- |
| Sham-W | 293.54 +/- 14.55 (22) | 299.45 +/- 9.60 (22) | 11.86 +/- 0.80 (22) |
| Sham-W/Na | 299.33 +/- 11.43 (24) | 294.46 +/- 9.57 (24) | 11.92 +/- 0.80 (24) |
| PAL-W/Na | 264.60 +/- 2.53 (15) | 265.60 +/- 2.80 (15) | 12.15 +/- 0.54 (15) |

Values are expressed as means +/- SE.

**Figure 4:**
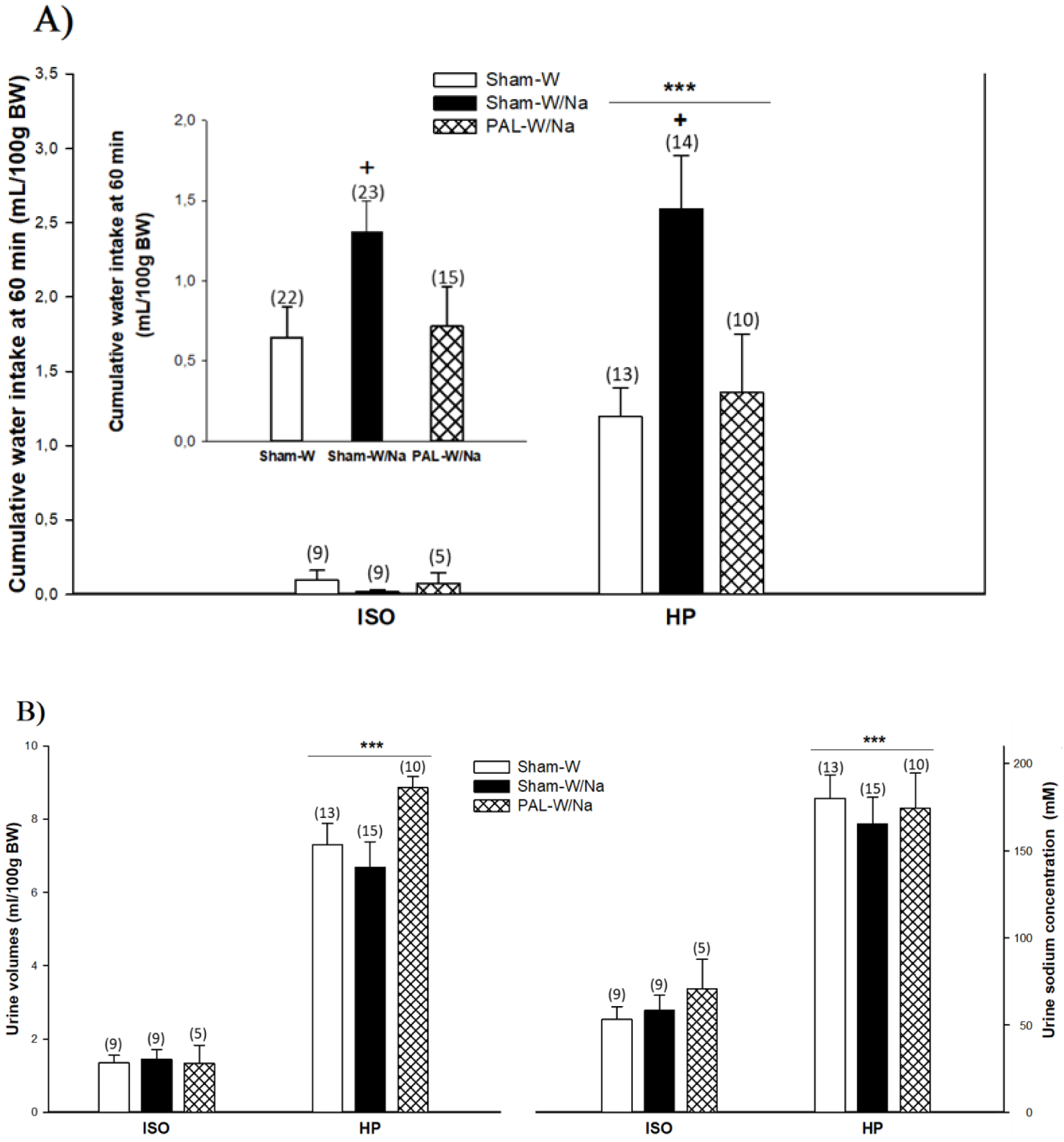
**A)** Cumulative water intake (mL/100g BW) during the 20-min iv infusion (0.15 mL/min) of a hypertonic (1.5M NaCl, HP) or isotonic (0.15M NaCl, ISO) solution and the following 60 minutes in adult offspring of each PM model. **B)** Urine volumes (mL/100g BW) and sodium concentration (mM) after the ISO or HP iv infusion in adult offspring of each PM model. Values are expressed as means +/- SE, (n). (^***^) Significant differences between ISO and HP infused animals, p < 0.0001. (+) Significant differences between Sham-W/Na with respect to Sham-W and PAL-W/Na groups, p < 0.05. ISO: isotonic sodium solution; HP: hypertonic sodium solution.

Relative to urine analysis, data indicated a significant increment in urine volumes and urine sodium concentration in hypertonic-infused animals relative to those infused with the isotonic solution [F (1, 52) = 154.94, p < 0.0001, η2p = 0.75, and F (1, 52) = 70.62, p < 0.0001, η2p = 0.58, respectively]. Data are shown in (**Figure 4B**). There was no significant main effect for PM or interaction between this factor and solution infused.

### 3.4 Experiment #4: Long-term PM effects in adult offspring challenged by a hypovolemic thirst model

A 3-way ANOVA was conducted with PM model and Furosemide-treatment as independent variables and both solutions (water and hypertonic sodium solution) as repeated measures. As expected, sodium-depleted animals drank greater volumes of fluids than non-depleted animals [Furosemide treatment-main effect: F (1, 68) = 204.11, p < 0.0001, η2p = 0.75]. Furthermore, animals drank significantly higher volumes of water than hypertonic sodium solution, independently of PM models and Furosemide-treatment received [Solution-main effect: F (1, 68) = 6.25, p = 0.0148, η2p = 0.08]. Regarding the PM model, a main effect of this factor [F (2, 68) = 8.48, p = 0.0005, η2p = 0.20] and two 2-way interactions PM model x Furosemide treatment and PM model x solution also achieved significance [F (2, 68) = 5.94, p = 0.0042, η2p = 0.15, and F (2, 68) = 5.42, p = 0.0065, η2p = 0.14, respectively].

Concerning the first 2-way interaction (PM x Furosemide), post hoc comparisons indicated that depleted-adult offspring of both PM models consumed lower volumes of fluids than Sham-W animals (**Figure 5B**). However, it is interesting to highlight that Sham-W/Na animals exhibited a stronger reduction in volumes consumed than PAL-W/Na group, leaving this last group with a middle consumption level between Sham-W/Na and Sham-W groups. Non-significant differences were found in fluid intake as a function of PM models in non-depleted animals (**Figure 5A**).

**Figure 5:**
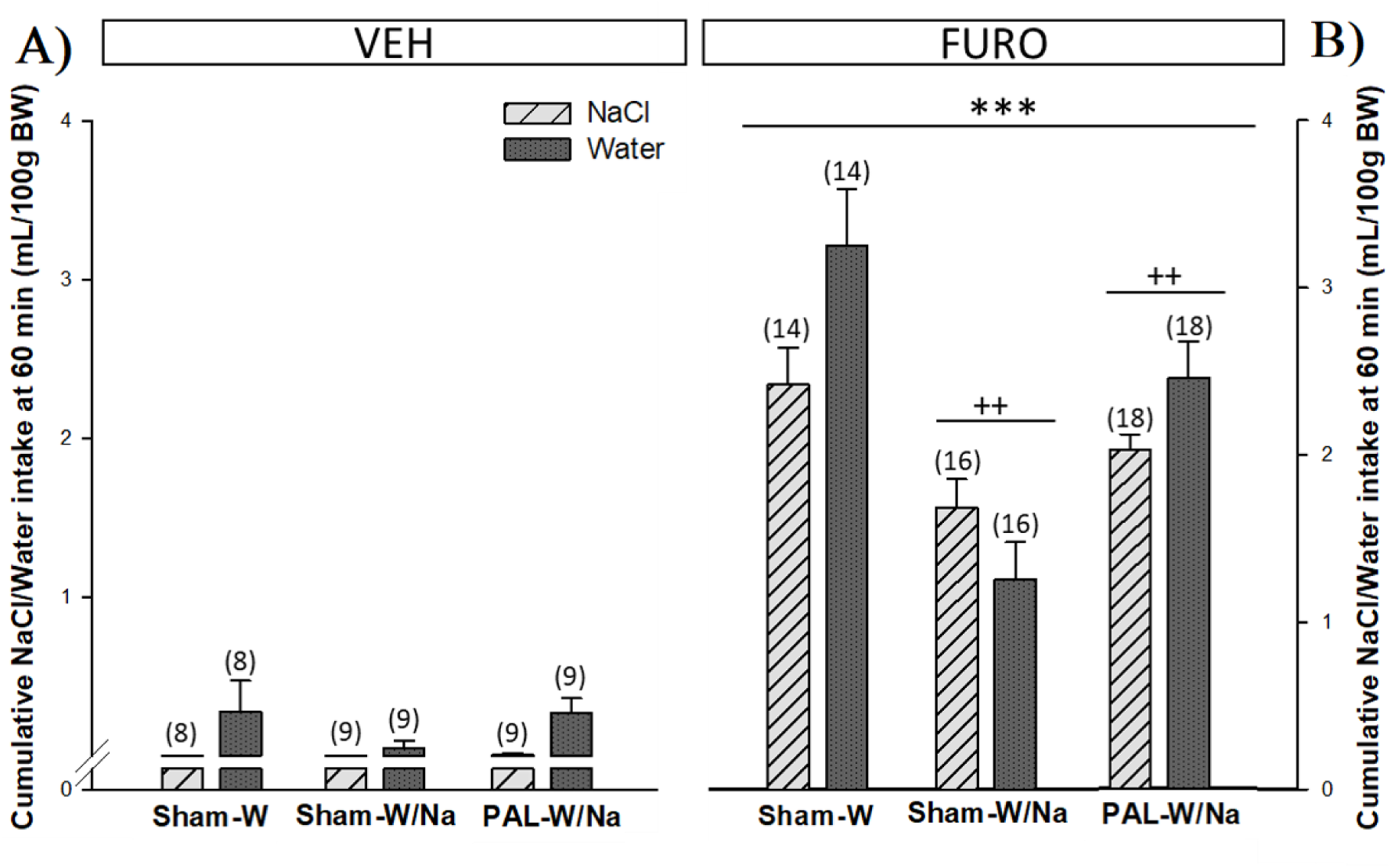
Cumulative water and 0.45M NaCl (mL/100g BW) at 60 min in adult offspring of each PM models in non-sodium depleted, VEH **(A)** and sodium-depleted FURO **(B)** conditions. Values are expressed as means +/- SE, (n). (^***^) Significant differences between VEH and FURO groups, p < 0.0001. (++) Significant differences between PM groups.

Regarding the 2-way interaction involving PM x solution, LSD test indicated a lower water intake in Sham-W/Na group while PAL-W/Na group exhibited, once again, a middle consumption level significantly different of both, Sham-W/Na and Sham-W groups. In relation to the hypertonic sodium intake, post hoc test revealed a significant reduction in the consumption levels of this solution in Sham-W/Na animals compared to the Sham-W group. The hypertonic sodium solution intake in the PAL-W/Na groups did not achieve statistical significance with respect to the other PM groups.

### 3.5 Experiment #5: Long-term PM effects on brain activation patterns (Fos + cells) in adult offspring exposed to a hypovolemic thirst model

In order to determine the PM effects on activation patterns in the LT, hypothalamus or brainstem in response to a hypovolemic thirst model, total number of Fos + cells was quantified in Furosemide depleted-adult offspring. Analyses were conducted via 1-way ANOVAs.

#### 3.5.1 Lamina terminalis nuclei

A significant effect of PM was found in the SFO [F (2, 10) = 11.72, p = 0.0024, η2p = 0.70]. A posteriori comparisons indicated a significant reduction in the number of Fos + cells in the SFO in adult offspring of both PM models. However, while the major reduction in the number of Fos + cells was observed in the Sham-W/Na group, the PAL-W/Na group exhibited a middle activation level between the Sham-W and Sham-W/Na groups (**Figure 6**). A correspondence was observed between the SFO’s activation levels and the consumed volumes of fluids in the test after the Furosemide depletion. Non-significant differences were detected in the activation patterns of MnPO and OVLT.

**Figure 6:**
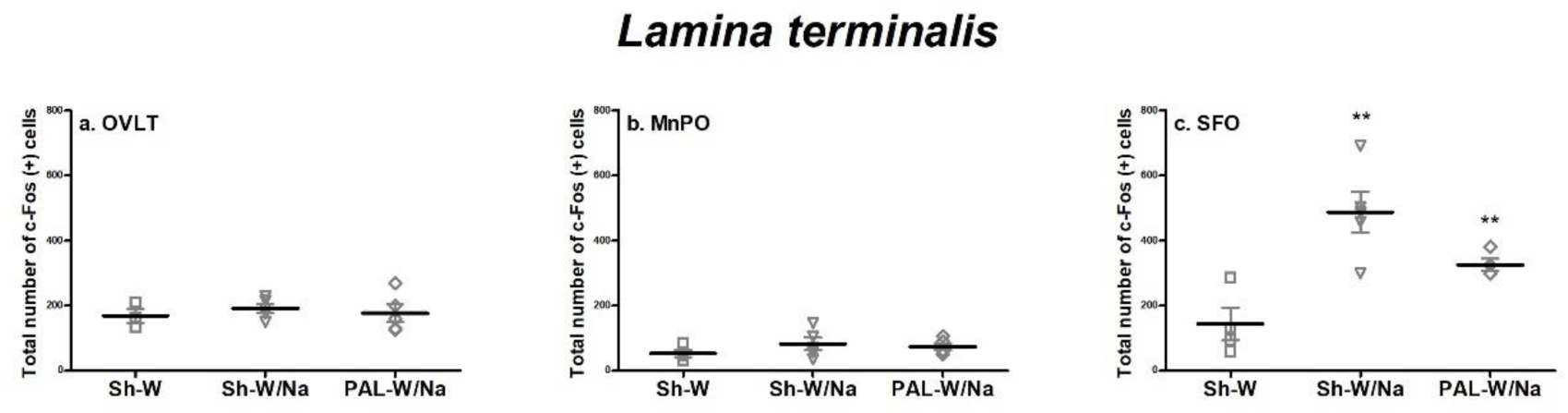
Number of Fos + cells in the *lamina terminalis* nuclei: OVLT (a), MnPO (b) and SFO (c). Values are expressed as means +/- SE. (^**^) Significant differences relative to the other groups, p < 0.001. **OVLT**: *organum vasculosum of the lamina terminalis*; **MnPO**: median preoptic nucleus; **SFO**: subfornical organ.

#### 3.5.2 Hypothalamic SON and PVN nuclei

A significant effect of PM was found in the SON, PaV and PaPo [F (2, 8) = 7.58, p = 0.0142, η2p = 0.65, F (2, 10) = 12.22, p = 0.0021, η2p = 0.71, and F (2, 9) = 4.61, p = 0.0418, η2p = 0.51, respectively]. When considering the SON, LSD test indicated a significant increases in the number of Fos + cells in Sham-W/Na groups, result that it is in accordance with those reported in Macchione et al., 2012. Furthermore, post hoc comparisons revealed a significant increase in the number of Fos + cells in the PaV in adult offspring of both PM models with respect to the Sham-W group. Once again, PAL-W/Na group expressed an intermediate activation level between Sham-W and Sham-W/Na groups, while the major increase in the number of Fos + cells relative to the Sham-W group was observed in the Sham-W/Na group (**Figure 7**). In the case of the PaPo nucleus, a significant increment in the number of Fos + cells was observed in both PM groups in relation to the Sham-W group. No significant differences were detected in PaLM, PaMM, and PaPM as a function of the PM models.

**Figure 7:**
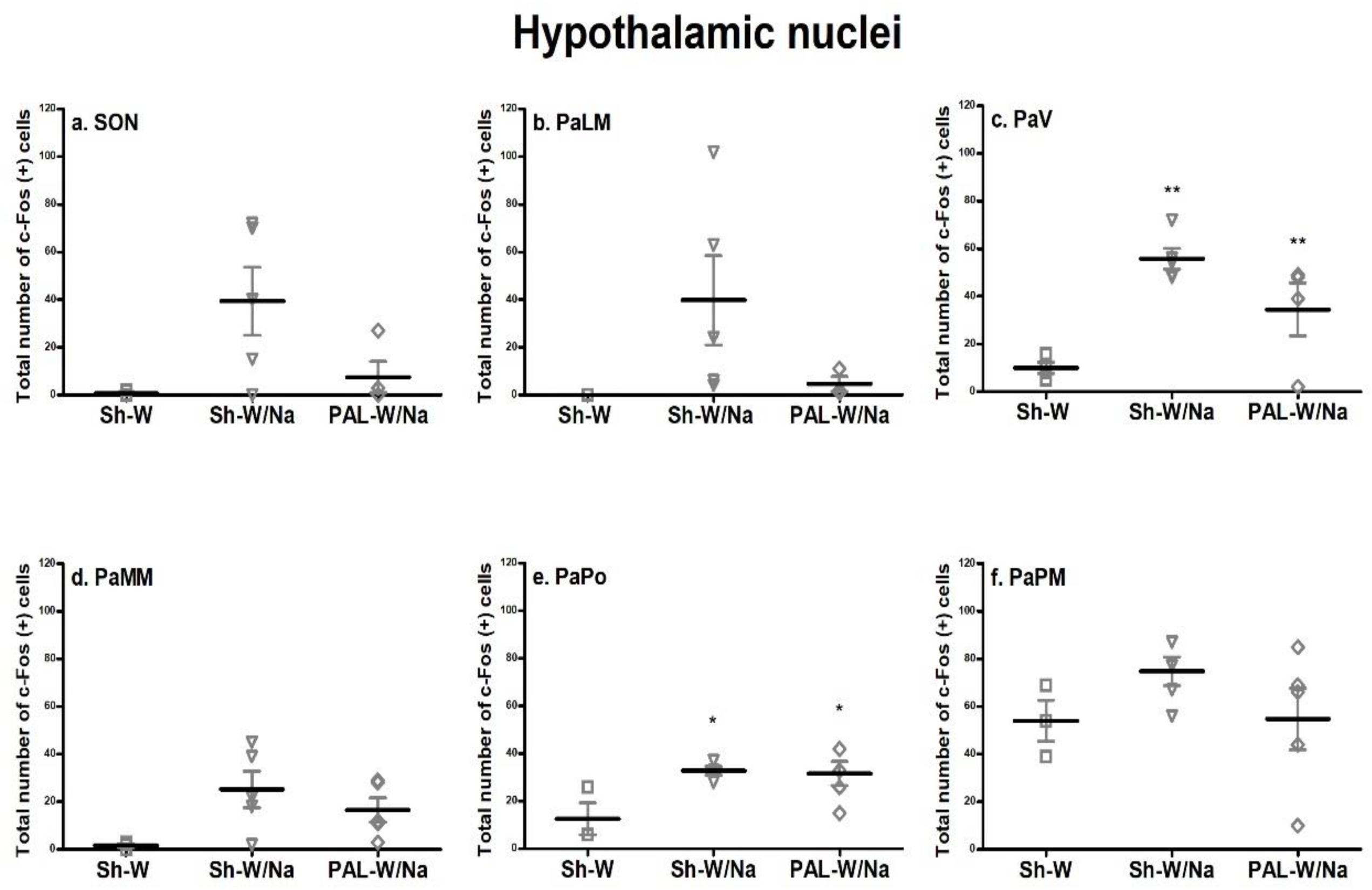
Number of Fos + cells in the follow hypothalamic nuclei: SON (a), PaLM (b), PaV (c), PaMM (d), PaPo (e) and PaPM (f). Values are expressed as means +/- SE. (^**^) Significant differences relative to the other groups, p < 0.001. (^*^) Significant differences between Sham-W/Na and PAL-W/Na groups with respect to Sham-W group, p < 0.05. **SON**: supraoptic nucleus; **PaLM**: paraventricular nucleus, lateral magnocellular subdivision; **PaV**: paraventricular nucleus, ventral subdivision; **PaMM**: paraventricular nucleus, medial magnocellular subdivision; **PaPo**: paraventricular nucleus, posterior parvocellular subdivision; **PaPM**: paraventricular nucleus, medial parvocellular subdivision.

#### 3.5.3 Brainstem nuclei

A significant effect of PM was found in the ventrodorsal subdivision of the DRN [F (2, 9) = 16.27, p = 0.0010, η2p = 0.78]. A posteriori comparisons indicated a significant increase in the number of Fos + cells in PAL-W/Na offspring with respect to Sham-W and Sham-W/Na groups **(Figure 8A & B)**. Considering the NTS, a reduction in the number of Fos + cells was observed in the PAL-W/Na and Sham-W/Na groups with respect to Sham-W group, although it did not achieve statistically significant levels [F (2, 7) = 3.72, p = 0.0793]. Neither significant differences nor interactions achieved significance in AP, LC, LPB, and DRN subdivision VL.

**Figure 8:**
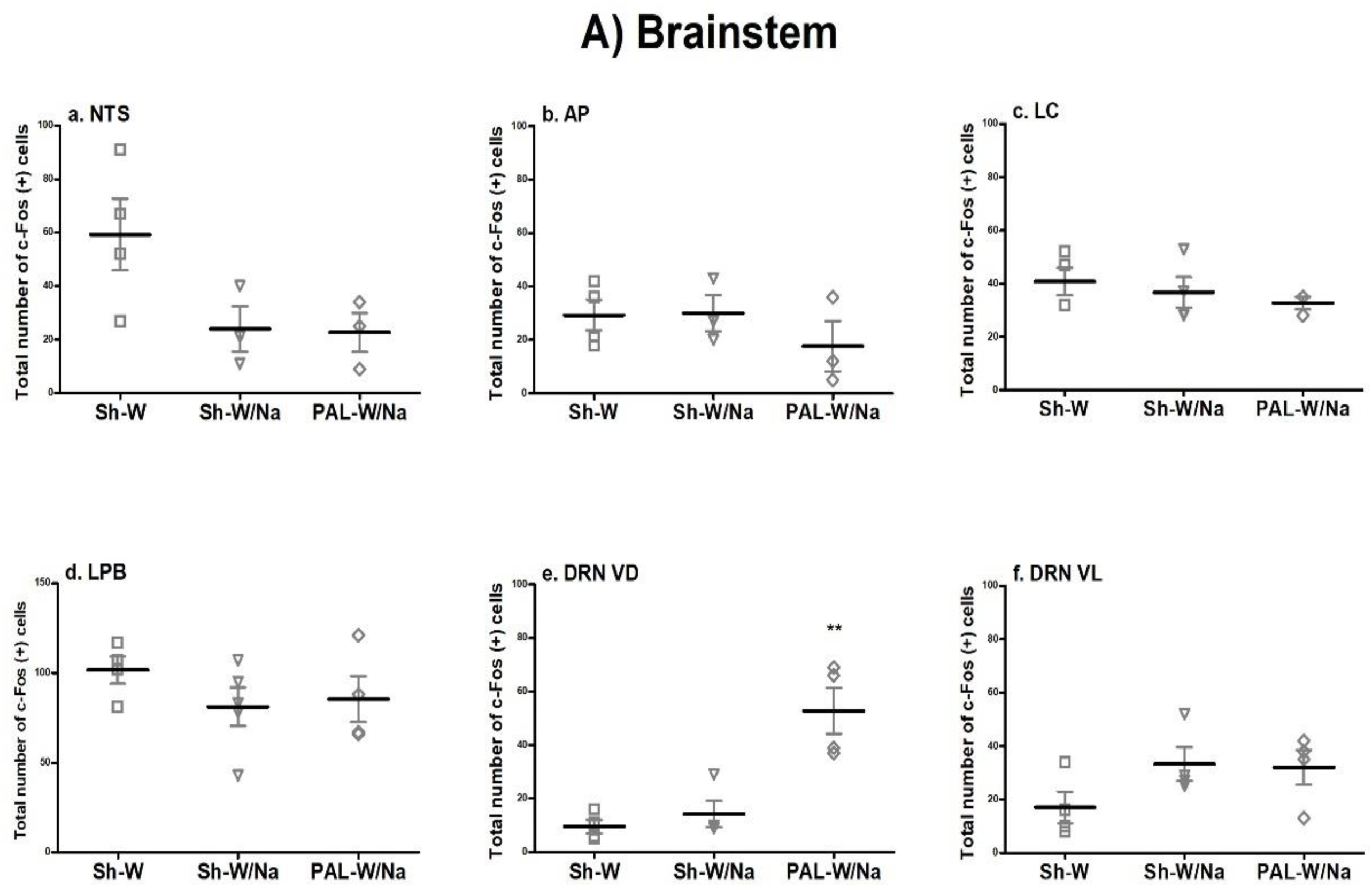
**A)** Number of Fos + cells in the follow brainstem nuclei: NTS (a), AP (b), LC (c), LPB (d), DRN VD (e) and DRN VL (f). Values are expressed as means +/- SE. (^**^) Significant differences between PAL-W/Na group with respect to Sham-W and Sham-W/Na groups, p < 0.001. **NTS**: nucleus of the solitary tract; **AP**: area postrema; **LC**: locus coeruleus; **LPB**: lateral parabrachial nucleus; **DRN VD**: dorsal raphe nucleus, ventrodorsal subdivision; **DRN VL**: dorsal raphe nucleus, ventrolateral subdivision.

**Figure 8:**
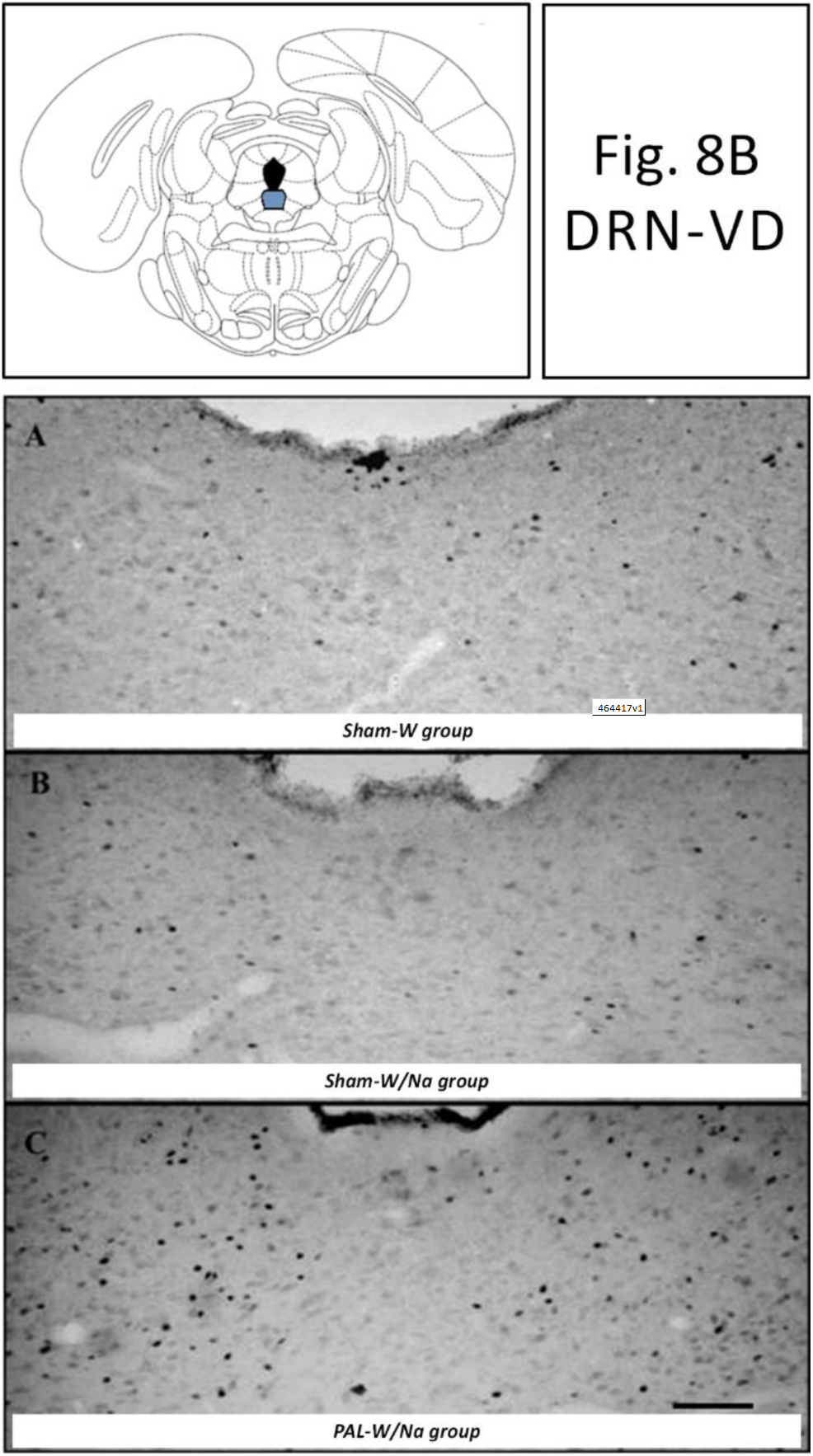
**B)** Representative photomicrographs showing the pattern of c-Fos immunoreactivity in the DRN VD of Sham-W (A), Sham-W/Na (B) and PAL-W/Na (C) after the Furosemide sodium depletion. In the upper square is represented the approximate antero-posterior level (to Bregma) where select brain regions were analyzed, based on Paxinos & Watson (2007). Magnification (10x). Scale bar = 100 m. **DRN VD**: dorsal raphe nucleus, ventrodorsal subdivision.

## 4. Discussion

Our results provide novel information about the different homeostatic systems and specific activation pattern of brain nuclei programmed by perinatal environmental cues which result in differential dipsogenic responses even during the adulthood. We found pieces of evidence that, depending on the characteristics of the perinatal environment that programmed the organism during the development; adult offspring expresses specific drinking responses which can be predicted according to the match between the osmotic challenge that triggers the response in the adult and the stimulus/stimuli that perinatally programmed this individual during its development.

Analyzing the key stimuli that can perinatally program the offspring in the PM models here employed, we focused on **Table 2** that shows the plasma parameters in 21-day-old female pups. Our results indicated that maternal free access to a hypertonic sodium solution significantly increased circulating sodium levels in weaning pups of both PM groups (Sham-W/Na and PAL-W/Na groups). However, this sodium increment was stronger in PAL-W/Na group and was also coupled with a significant increase in osmolality levels. This hypernatremia in the offspring, induced by maternal high salt intake, was previously reported by Deloof et al., 2000; Jelinek et al., 1990; Macchione et al., 2012. It is important to note that, the hypernatremia and hyperosmolality observed in weaned pups’ plasma parameters were not found in dams. Diverse causes can be considered to explain this apparent discrepancy between weaned pups and maternal values. One of them is that renal development and capacity of excretion in weaning pups may not be enough to eliminate the excess of sodium that comes from amniotic fluid, milk and/or the hypertonic sodium solution directly consumed by the offspring during the last week of lactation. Maternal natriophilia has been reported to result in an electrolyte accumulation in pups’ body because diuretic and sodium efficiency is lower at this age (Deloof et al., 2000; Misanko et al., 1979). High maternal salt intake influences fetal kidney development, inducing the adult offspring to be more susceptible to salt retention, probably increasing blood pressure later in life (Gray et al., 2013). Parameters such as days of gestation and number of pups/litter, did not show differences as a function of the PM models suggesting that, also the PAL-W/Na model, did not significantly alter the normal development of the offspring, in accordance with previous reports (Argüelles et al., 2000; Perillan et al., 2004).

PAL-W/Na dams had an exacerbated water and total fluids intake relative to sham-ligation dams, as it was reported by Costales et al., 1984; J. Fitzsimons & Simons, 1969. An evident natriophilia was also observed in PAL-W/Na dams (more than 20 mL/day) although it did not reach significant differences with respect to Sham-W/Na dams. Hence, we emphasize that this type of experimental paradigm in which pregnant animals are exposed to a rich source of sodium highlights the clear increment in voluntary sodium consumption from the beginning of the gestational period. In our records, Sham-W/Na and PAL-W/Na dams consumed an accumulative hypertonic sodium solution of 836.80 +/- 115.22 mL and 1,202.00 +/- 130.98 mL, respectively. In perspective, we must take into consideration that it normally occurs in nature and especially in occidental society with the strong impact that it can have on the health of the mother and the baby.

PAL also modified kidney weights. An ischemia and reduction of approximately 50% of the kidney subject to the partial ligation was observed as a consequence of the decrease in the renal blood flow. This renal blood flow reduction triggers a chronic release of renin from the juxtaglomerular apparatus. The ischemia of the ligated-kidney was compensated by a hypertrophy of the opposite kidney to supply the renal function. In accordance with previous research (Perillan et al., 2004; Vijande et al., 1996), we corroborated that PAL elicited a chronic over-activation of RAS, through the quantification of the maternal PRA levels. Manipulations on maternal blood pressure and PRA levels caused equivalent modifications on fetal blood pressure and PRA (Lumbers & Lewes, 1979).

A significant decrease in osmolality with normal levels of plasma [Na+] was observed in PAL-W/Na dams at weaning. We suspect that the predictable increase in natremia in this PM model was not observed in our quantifications because the intake episodes normally occur overnight, and we collected samples at the midday. In adults, natremia quickly return to their normal levels because the fully mature homeostatic systems. Furthermore, in our model the hypertonic solution intake was voluntary, intermittent and it was accompanied by water intake; unlike other protocols in which animals have a mandatory access to the rich source of sodium through diet or through a hypertonic solution in replacement of water (Deloof et al., 2000; Lowe et al., 1992). Experimental conditions here employed allowed the dams to drink an isotonic cocktail between water and sodium, satisfying their sodium appetite but maintaining the hydrosaline balance. The same hypothesis is also valid to explain why there were no changes in plasma sodium levels in Sham-W/Na dams. On the other hand, PAL through endocrinal (RAS over-activation) and behavioral (polydipsia) responses, induced a fall in the osmolality in PAL-W/Na dams. This alteration in osmolality may suggest a volume expansion (Andrea Godino et al., 2013) due to a false signal of blood flow drop conducting to drink larger volumes of fluid, predominantly water. Probably, long-term effects of the chronic activation of RAS may be the cause of the decreased plasma osmolality observed in dams at the weaning.

When the offspring were defied with an absolute dehydration model (overnight water deprivation), PAL-W/Na offspring responded similarly to Sham-W/Na and both programmed groups drank low volumes of water with respect to non-programmed (Sham-W) descendants. The programmed offspring were less responsive to the dehydration state, suggesting an upward resetting of the osmotic threshold for onset of thirst, probably due to a desensitization of the angiotensin II type 1 (AT1) receptor. Several studies have established that perinatal hyperosmolality and/or hypernatremic environment elicits an osmotic threshold shift or a reset central osmostat in the adult offspring (Gray et al., 2013; Ross et al., 2005). The reset of the osmostat can be evidenced, for example, by a decrease in water intake against certain stimuli that trigger thirst, such as circulating ANG II. Typically, many G-protein–coupled receptors, like AT1 receptor, are desensitized after repeated ligand exposure, so repeated ANG II abolishes calcium responses to subsequent ANG II (Gebke et al., 1998; Santollo et al., 2018; Thomas et al., 1996). Furthermore, models that elicit thirst by injections of ANG II have demonstrated that the AT1 desensitization reduced drinking response after pretreatments of ANG II (Vento et al., 2012; Vento & Daniels, 2014).

Programmed offspring also exhibited polyuria (excreting high urine volumes) after the intake test. While non-programmed offspring excreted low volumes of urine independently the amount of water consumed (r = -.08, p = .98), Pearson’s analysis indicated a positive correlation between water volumes consumed and urine volumes excreted in programmed animals that reached significant levels in the case of PAL-offspring (Sham-W/Na: r = .54, p = .07, and PAL-W/Na: r = .70, p = .02). In other words, the programmed animals rapidly excreted what they drank rather than retain the water consumed, as the non-programmed seem to do. These observations together showed long-term programming effects on the homeostatic balance, probably determining a particular balance status in programmed offspring. Water deprivation increments osmolality and decreases plasma volume by inducing homeostatic responses that include thirst, RAS activation and increases of ANG II circulating to restore the body’s water balance (Da Silveira et al., 2007; Gottlieb et al., 2006; Greenwood et al., 2015; Johnson & Thunhorst, 2007; Stricker & Sved, 2000). However, these conditions also characterized the PM conditions studied here (i.e. hyperosmolality and RAS activation) supporting the hypothesis that programmed offspring should respond similarly to each other and be less thirsty that non-programmed animals, probably due to a desensitization in the switch-on of the circuits or central centers triggered by intra- and extracellular dehydration.

We also analyzed a thirst model induced by hypertonic sodium chloride infusion (relative hyperosmotic thirst), where the main stimuli are the hyperosmolality and hypervolemia activating central and peripheral osmo-sodium receptors, and volume/arterial baroreceptors but not the peripheral RAS. Taking these conditions into account, when the adult offspring was challenged with a sodium overload, we found that PAL-W/Na offspring responded like the control Sham-W group consuming similar water volumes at the test. This behavior may be due to the fact that the main programmer stimulus, the peripheral RAS over-activation, is not present in this thirst model. On the other hand, drinking differences were only observed in the Sham-W/Na offspring, with a higher water intake after 60-min of the test, as previously reported in Macchione et al., 2015. We postulate that sodium overload employed to elicit thirst (iv hypertonic sodium infusion) reveals significant differences only in the Sham-W/Na group because both environmental conditions (perinatal- and test-conditions) match with each other and, thus, affect differently the baroreceptors reactivity and thirst.

Concerning the long-term PM effects on adult sodium-depleted offspring, our results revealed i) PAL-W/Na drank higher volumes of both fluids (water and hypertonic sodium solution) relative to Sham-W/Na offspring but, ii) both programmed groups drank lower volumes with respect to non-programmed animals (Sham-W group). Previous works that studied PAL programming model employed, as a PM control group, one which we have established in this work as a Sham-W/Na group. In those studies, authors reported that PAL programming effects incremented fluid induced-intake on adult offspring compared to their control group (here Sham-W/Na) (Argüelles et al., 2000; Vijande et al., 1996). When considering Sham-W/Na as the control group, the interpretation of that, PAL exacerbated thirst and sodium appetite on adult offspring after a sodium depletion, is absolutely correct. Although, if it is taking into account that perinatal sodium availability is a programming stimulus itself, and an additional control group is conducted (without sodium availability, here Sham-W), a new interpretation of the results should be considered. In comparison to this new control group, we demonstrated that both programmed offspring were lower drinkers. These long-term changes in drinking behavior can be initially attributed to both perinatal events, such as maternal hormonal alterations (high maternal PRA levels) in PAL-W/Na group, and early hydroelectrolyte changes (hypernatremia) in both programmed-groups. This last condition was observed in the breast milk of PAL mothers (Vijande et al., 1996) or in plasma sodium and osmolality of female pups, as reported here. It is well established that Furosemide sodium-depletion induces a hypovolemic/hyponatremic status that stimulates central and peripheral RAS (Fluharty & Epstein, 1983; Fregly & Rowland, 1985; Rowland & Morian, 1999). Due to the RAS activation, animals consume high volumes of sodium and water, restoring the body sodium status (Fitzsimons, 1998; Rowland & Morian, 1999). While Sham-W/Na’s drinking patterns obey the perinatal hypertonic sodium availability that programmed a new set point to restore the fluid balance resulting in a hypertonic balance status (Contreras and Ryan, 1990; Curtis et al., 2004; Macchione et al., 2012), PAL-W/Na offspring were programmed by a mixture of stimuli (i.e. sodium overload plus chronic RAS over-activation). This combination of stimuli also elicits a low-drinking phenotype by a new set point of thirst to reestablish body fluid status. Once again, these results are in agreement with our hypothesis suggesting that when the osmotic challenge matches with the perinatal programming stimuli, both programmed groups respond differently with respect to the non-programmed group.

The activation patterns in the SFO may explain, at least in part, this “low-thirsty” phenotype drinking profile found in both programmed groups when were challenged by the hypovolemic thirst model. On one hand, as we previously reported in Macchione et al., 2012, sodium programmed animals (Sham-W/Na) drank low fluid volumes and exhibited an increased number of Fos + cells in the SFO, in comparisons with non-programmed animals (Sham-W group). SFO contains cells that are sensitive to humoral signals, such as osmolality, sodium concentration, and circulating ANG II levels in plasma/CSF because of its neurons contain osmo-sodium- and angiotensinergic receptors (Gutman et al., 1988; Krause et al., 2008; Noda, 2006). Therefore, it seems likely that during the PM period, the excitability of the SFO caused by the increase in circulating sodium levels could have triggered a sensitization of the SFO, explaining the increment in the number of Fos + cells observed in the Sham-W/Na group. On other hand, PAL-W/Na also exhibited high number of Fos + cells in the SFO coupled with a lower water intake at the sodium-depletion test. In this PM group, we also found a significant increase in the number of Fos + cells at the DRN-VD. We postulate that the DRN-VD activation pattern may suggest a failure in the disinhibition of the serotonergic system allowing the high level of SFO activation. Studies carried out by our and other laboratories have provided numerous evidence that the DRN forms a circuit that remains active during the hydroelectrolyte balance, exerting a tonic inhibition on water and sodium intake (Badauê-Passos et al., 2007; Cavalcante-Lima et al., 2005b, 2005a; Franchini et al., 2002; Olivares et al., 2003). The expected response to a homeostatic state that requires the intake of both water and sodium is a deactivation of this nucleus. It has been shown that the activity of serotonergic cells in the DRN is affected by the body sodium status, decreasing its activity during a sodium depletion (Cavalcante-Lima et al., 2005b, 2005a; Franchini et al., 2002; A Godino et al., 2007; Olivares et al., 2003). However, in the case of PAL-W/Na model, our data failed to observe this deactivation, suggesting that the chronic RAS stimulation during the perinatal period would have desensitized this circuit. As mentioned before, sodium-depletion stimulates central and peripheral RAS (Fluharty & Epstein, 1983; Fregly & Rowland, 1985; Rowland & Morian, 1999). This group of animals appears to be more resistant to the expected deactivation when an acute sodium depletion occurs. Contrasting the neuronal activation observed in the DRN with the patterns of induced-water and sodium intake during the test, we can notice a relationship between both events. In relation to the non-programmed animals, PAL-W/Na offspring showed a higher number of Fos + cells in the ventrodorsal subdivision of the DRN after the Furosemide sodium depletion, while it exhibited a reduced fluid intake.

The modulation of water and sodium intake involves the interaction between the LT and hindbrain serotonergic inhibitory systems (Badauê-Passos et al., 2007; Franchini et al., 2002; A Godino et al., 2007, 2010; Tanaka et al., 2004). Indeed, it has been determined a bidirectional angiotensinergic neural pathway between the SFO and DRN (Lind, 1986) and that the LT neurons activated by a F/SLD treatment are connected with the DRN (Badauê-Passos et al., 2007). The administration of ANG II into the SFO inhibits the activity of the DRN neurons and, related to these observations, the activation of the SFO-DRN angiotensinergic pathway induces a local reduction in the release of serotonin into the DRN (Tanaka et al., 1998, 2003). These evidences suggest that neurons projecting from the SFO to the DRN are monitoring the levels of circulating ANG II and sending this information to the DRN. Taking into account the implications of the DRN in fluid intake regulation, we propose that the high levels of ANG II circulating chronically during the perinatal period in PAL-W/Na offspring could desensitize this angiotensinergic SFO-DRN pathway.

In summary, the results reported here support the hypothesis that perinatal programming is a phenomenon that differentially affects particular systems and target sites, inducing specific dipsogenic responses depending on the coincidence between perinatal programming conditions and the osmotic challenge in the latter environment. Regarding this, PAL-offspring responded similarly to early sodium exposed animals when were challenged with osmotic conditions similar to those that acted as program stimuli during the perinatal period (i.e. persistent hypernatremia and peripheral RAS activation), like overnight water deprivation and Furosemide sodium depletion. However, when adult offspring were defied with a thirst model that mainly involves the osmo-sodium-sensors and excludes the RAS system (relative dehydration induced by a hypertonic NaCl intravenous infusion), PAL/W-Na group responded as a non-programmed group because the challenged conditions did not entirely match with the perinatal programming conditions.

## Acknowledgments

This work was supported by grants from the Agencia Nacional de Promoción Científica y Tecnológica (ANPCyT, PICT 2016 N° 0869), PUE-CONICET (RES. N°2555/16) and Secretaría de Ciencia y Tecnología (SECyT, RES. N° 411/18). FMD holds a postdoctoral fellowship from the Consejo Nacional de Investigaciones Científicas y Técnicas (CONICET). Furthermore, the authors thank Micaela A. Macchione for polishing the language of the manuscript.

